# Disruption of hemocyte differentiation and distribution in Drosophila Ptr^23c^ mutants

**DOI:** 10.1101/2025.07.03.662979

**Authors:** Cristina Parada, María Constanza Silvera, Tabaré de los Campos, Rafael Cantera, Carmen Bolatto, Daniel Prieto

## Abstract

In *Drosophila*, hemocytes are essential for development and immunity, with their differentiation and spatial distribution under strict regulation. Here, we examine the effect of the *Ptr*^*23c*^ null mutation in embryonic hemocyte development. Contrary to our initial hypothesis, *Ptr* does not regulate hemocyte number, as the mutation did not affect the total number of hemocytes, apoptosis, mitosis, or the balance between major subpopulations. However, *Ptr*^*23c*^ mutants displayed disrupted distribution and premature hemocyte differentiation, marked by accelerated maturation at stage 12. Despite this early differentiation, *Ptr*^*23c*^ embryos exhibited a 50% reduction in mature hemocytes by stage 16, as quantified by *serpent*-driven mCherry expression. Our findings establish Ptr as a regulator of hemocyte distribution and differentiation timing during normal embryogenesis, possibly through modulation of the *serpent* pathway.

## Introduction

Patched-related (Ptr) is a conserved transmembrane protein belonging to the patched domain-containing protein family (Kuwabara and Labouesse, 2002; Pastenes et al., 2008). In *Drosophila melanogaster*, Ptr can interact directly with the signaling protein Hedgehog (Hh) and *Ptr*^*23c*^ null mutants exhibit phenotypic defects similar to those observed in Hh pathway mutants (Bolatto et al., 2022). Furthermore, cell culture assays suggest that Ptr acts as a negative regulator of Hh signaling (Bolatto et al., 2015, 2022).

*Ptr* is expressed in hemocytes (Bolatto et al., 2015; Cattenoz et al., 2020), which are arthropod blood cells, though its precise role in these cells remains unclear. Hemocytes are essential for immunity and development, with *Drosophila* possessing three main subtypes: plasmatocytes, crystal cells (CCs), and lamellocytes (Honti et al., 2014; Nappi and Vass, 1993; Yu et al., 2022). During *Drosophila* development plasmatocytes are, by far, the most common type of hemocyte. Constituting around 95% of circulating hemocytes, they function as phagocytes, bearing similarities to mammalian macrophages and monocytes (Gold and Brückner, 2015; Meister, 2004; Yoon et al., 2023). These migratory cells are responsible for phagocytizing apoptotic cells, tumor cells and invading microorganisms. They recycle the nutrients from the material that they have ingested and secrete antimicrobial peptides and components of the extracellular matrix (Evans and Wood, 2014; Krejčová et al., 2024; Wang et al., 2014).

Due to evolutionary conservation in hematopoiesis (Evans et al., 2003; Gold and Brückner, 2015; Lebestky et al., 2000), *Drosophila* hemocytes serve as a model for studying immune responses, tissue remodeling, wound healing, and even metastasis (McDonald and Montell, 2004; Yoon et al., 2023).

Embryonic hemocytes originate from procephalic mesodermal derivatives beginning from stage 8 onwards. At this stage, a population of mesodermal precursors undergoes four rounds of mitosis and reach a total of 600-700 cells (Tepass et al., 1994; Gyoergy et al., 2018). These numbers remain stable until the end of embryonic development (Vlisidou and Wood, 2015).

During germ-band extension hemocytes colonize the head region before migrating across the epithelial barrier between the head and tail (Tepass et al., 1994; de Velasco et al., 2006; Siekhaus et al., 2010). They then disperse along three stereotyped migratory paths: the ventral midline (dorsal and ventral to the developing ventral nerve cord), the intestinal primordium, and beneath the epidermis, along the developing dorsal vessel (Cho et al.; 2002 Ratheesh et al., 2015). As they migrate, hemocytes deposit extracellular matrix components and engulf cell debris resulting from developmental apoptosis (Tepass et al., 1994; Sonnenfeld and Jacobs, 1995; Manaka et al., 2004).

By stage 15 of embryogenesis, the hemocytes are evenly dispersed throughout the embryo, yet they still have the capacity to migrate to wound sites (Fauvarque and Williams, 2011; Moreira et al., 2011; Ratheesh et al., 2015; Tepass et al., 1994). Targeted ablation of hemocytes results in elevated embryonic lethality and defects in the ventral nerve cord (Defaye et al., 2009), as well as pupal lethality (Matsubayashi et al., 2017; Stephenson et al., 2022).

Our previous research revealed that late *Ptr*^*23c*^ embryos have reduced hatching ability (Bolatto et al., 2022; Parada and Prieto, 2025), which might explain their substantial embryonic lethality (Bolatto et al., 2015). Increased mortality was also observed at subsequent developmental stages (Parada and Prieto, 2025) and an apparent reduction in the number of embryonic hemocyte has been suggested (Bolatto et al., 2015). Given the fundamental role of hemocytes in embryonic development, a decrease in their number could potentially contribute to the reduced viability of *Ptr*^*23c*^ embryos and larvae. In this study, we investigate the effect of *Ptr*^*23c*^ mutation in the number, proliferation, apoptosis and distribution of hemocytes during embryogenesis. The specific mechanisms affected in *Ptr*^*23c*^ mutants remain unknown.

## Materials and Methods

### Genetics and Drosophila strains

*srpHemo-H2A::3xmCherry* was kindly gifted by Dr. Daria Siekhaus (Gyoergy et al., 2018). This stock was used to generate both the control strain (+*/+, twi-GFP*; *srpHemo-H2A::3xmCherry/srpHemo-H2A::3xmCherry*) and the *Ptr*^*23c*^*/CyO, twi-GFP*; *srpHemo-H2A::3xmCherry/srpHemo-H2A::3xmCherry* strain. Flies were maintained on a standard cornmeal, yeast and glucose medium at 25°C with 12h-dark and light cycles.

### Embryo collection and fixation

Flies were allowed to lay eggs on apple-juice agar plates supplemented with yeast. The eggs were collected, rinsed with water, dechorionated in 5% sodium hypochlorite and washed with PBST (0.05% Triton X-100 in phosphate-buffered saline). The embryos were then fixed by shaking them in a 1:1 mixture of heptane and 4% paraformaldehyde (PFA) for 20 minutes. The vitelline membrane was removed by replacing the PFA with cold methanol (−20°C) and shaking for 1 minute. The fixed embryos were then stored in fresh methanol at −20°C until further use.

### Immunofluorescence

The embryos were rehydrated and incubated with PBST 3% BSA for 1 hour. Immunostaining was performed using the following antibodies: rabbit anti-phosphorylated Histone H3 (1:200, Cell Signaling Technologies #9713), rat anti-mCherry (1:500, Invitrogen #M11217) or rabbit anti-Dcp-1 (cleaved *Drosophila* Death caspase-1) (1:100, Cell Signaling Technologies #9578). The embryos were then incubated with the antibodies at 4°C overnight, washed with PBST and incubated for 2 hours with the following fluorescently conjugated antibodies goat anti-rat (1:500, Invitrogen, A11011), goat anti-rabbit (1:500, Invitrogen #A11034) or Alexa Fluor 647-conjugated Peanut Agglutinin (PNA, 10 mg/ml, Molecular Probes #L-32460). After washing with PBS, the embryos were mounted in 80% Tris-buffered glycerol.

### Microscopy, Quantification and Statistical Analysis

Confocal images were acquired using a Zeiss LSM800 laser-scanning confocal microscope using either an EC Plan-Neofluar 20x/0.50 or a Plan-Apochromat 63x/1.40 M27 objective with GaAsP-PMT and Multialkali detectors. Images were analyzed using FIJI for Linux (Schindelin et al., 2012).

To calculate the mitotic index, stacks of confocal images taken along the longitudinal axis of vertically mounted embryos were processed and quantified using Object Counter 3D. Double-stained nuclei were identified in a new stack generated from the original channels in Image calculator using the logical operator ‘AND’, followed by Object Counter 3D particle analysis excluding sizes under 5 μm^3^.

To quantify the total number of hemocyte and apoptotic hemocytes, embryos were mounted between two coverslips and z-stacks of confocal images were obtained orthogonally to the longitudinal body axis were obtained for every half embryo. Whole embryos were reconstructed in three dimensions with FIJI, and quantification was performed using Object Counter 3D particle analysis.

A FIJI/ImageJ macro was developed specifically for analyzing hemocyte distribution. The perimeter of the embryo was determined using PNA staining and divided into 19 equally spaced concentric regions of interest (ROIs). We measured mCherry integrated density within each ROI. Each value was then expressed as a percentage of the total for each embryo. Data analysis was performed using either GraphPad Prism (version 6.01 for Windows, GraphPad Software, San Diego, CA, USA, www.graphpad.com) or R/Bioconductor 4.4.1.

### Flow Cytometry

A pool of 150 dechorionated embryos was disaggregated in HL solution (Salmand et al., 2011) using a Dounce homogenizer. The mixture was then filtered through a 100 μm mesh and centrifuged at 400 g for 5 min. The pellet was then resuspended in HL containing 2 mM EDTA and filtered through a 40 μm mesh. The cells were then fixed for 10 min in 4% PFA, washed in PBST, acquired using an Attune™ NxT cytometer, and analyzed in R/Bioconductor 4.4.1 based on a previously described method (Gonzalez et al., 2023). The fluorescence emitted by the mCherry reporter in hemocytes at three embryonic stages was quantified to obtain a discrete time series for the *Ptr*^*23c*^ strain and the control strain.

## Results

### Quantification of hemocyte number and death

Based on preliminary data, it was suggested that the *Ptr*^*23c*^ null mutation reduces the number of hemocytes in *Drosophila* embryos (Bolatto et al., 2015). To investigate this possibility, we introduced a hemocyte-specific nuclear mCherry reporter into the *Ptr*^*23c*^ strain to enable the identification of individual hemocytes (Gyorgey et al., 2018). Confocal microscopy stacks were used to count all hemocytes in each embryo examined at the onset of stage 16 (Figure 1A). In the control sample, we found the median count was around 600 cells (Figure 1B), which is consistent within the counts obtained using both the same methodology (Gyoergy et al., 2018) and a different methodology (Tepass et al., 1994). The same result was obtained when examining the *Ptr*^*23c*^ mutant samples. The median hemocyte count was 601 for control embryos (n = 15) and 618 for *Ptr*^*23c*^ embryos (n = 14) (Figure 1B).

**Figure 1.**
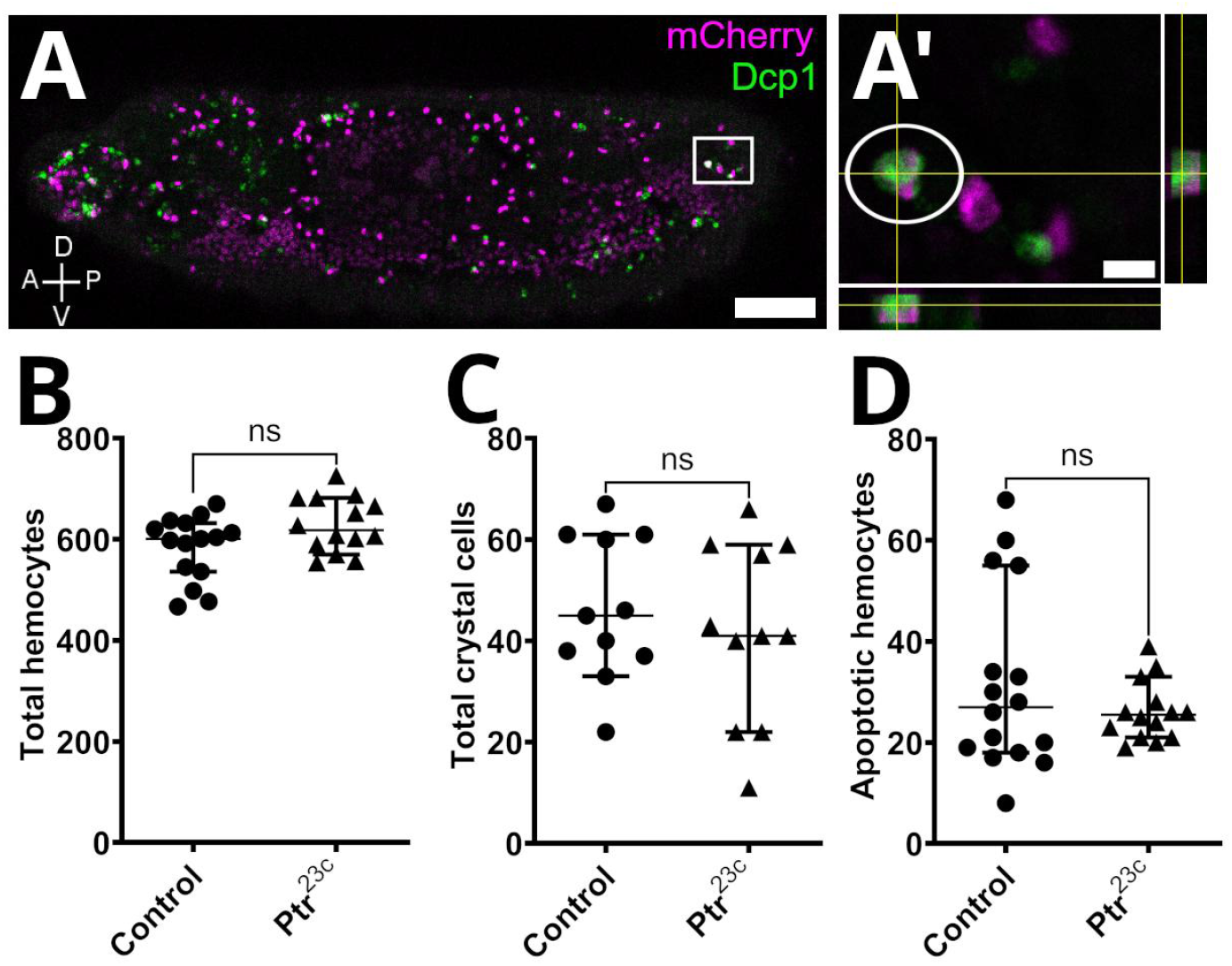
Hemocyte number, death, and distribution at stage 16 in *Ptr*^*23c*^ embryos. (A and A’) Embryonic hemocytes, which express the nuclear marker mCherry (*srpHemo-H2A::3XmCherry*), were immunostained using anti-Dcp-1 (cleaved *Drosophila* Death caspase 1). (A) Apoptotic hemocytes were defined by the colocalization of the apoptotic marker and the hemocyte-specific nuclear mCherry signal (white circle). (A’) Orthogonal optical planes are shown. The number of (B) total hemocytes, (C) total crystal cells and (D) apoptotic hemocytes were counted in embryos of each genotype. Scale bars: (A) 50 μm; (A’) 5 μm.

Having observed that the mutation did not appear to alter the total number of hemocytes present at stage 16, we investigated whether it caused a shift in the relative proportions of the two major hemocyte populations in the embryo, plasmatocytes and CCs. At stage 16, CCs can easily identified from other hemocytes because they remain clustered around the proventriculus and possess large nuclei (Lebestky et al., 2000). Interestingly, we observed no significant difference in the number of CCs in *Ptr*^*23c*^ embryos (median = 41, n = 11) compared to controls (median = 45, n = 11) (Figure 1C).

In order to investigate whether the *Ptr*^*23c*^ mutation increases apoptosis among embryonic hemocytes, we counted the number of particles that were positive for both the hemocyte marker *srp*Hemo-H2A::3XmCherry and the apoptotic marker Dcp-1 (see Figure 1A and A’). Stage 16 was chosen for this analysis since *Ptr* expression in hemocytes of a wild-type strain has been demonstrated at this stage (Bolatto et al., 2015; Cattenoz et al., 2020), and our initial observation of an apparent reduction in *Ptr*^*23c*^ hemocyte numbers was made at this stage (Bolatto et al., 2015). No differences were detected in the number of apoptotic hemocytes (*Ptr*^*23c*^ median = 25.5, n = 14; control median = 27, n = 16; see Figure 1D) or in total apoptosis (data not shown).

### Hemocyte distribution

As the *Ptr*^*23c*^ mutation did not affect the total number of hemocytes, we investigated whether an altered distribution pattern at stage 16 could explain the previous observation of fewer number of hemocytes in the ventral region of *Ptr*^*23c*^ embryos (Bolatto et al., 2015). To this end we developed a FIJI/ImageJ macro that defined concentric bands as regions of interest (ROIs) ranging from the periphery (ring 1) to the center (ring 19) of the embryo (n=15) (see Figure 2A, left panel). Notably, we found a significant difference in hemocyte distribution between *Ptr*^*23c*^ and control embryos (W= 53371, p= 8.6×10^−11^, Wilcoxon rank sum test with continuity correction) (Figure 2B). Hemocytes were less abundant at the periphery and center of the *Ptr*^*23c*^ embryos relative than in controls, with significant differences (p < 0.05, Wilcoxon U-test) observed in rings 1, 2, 17, 18, and 19 (Figure 2B).

**Figure 2.**
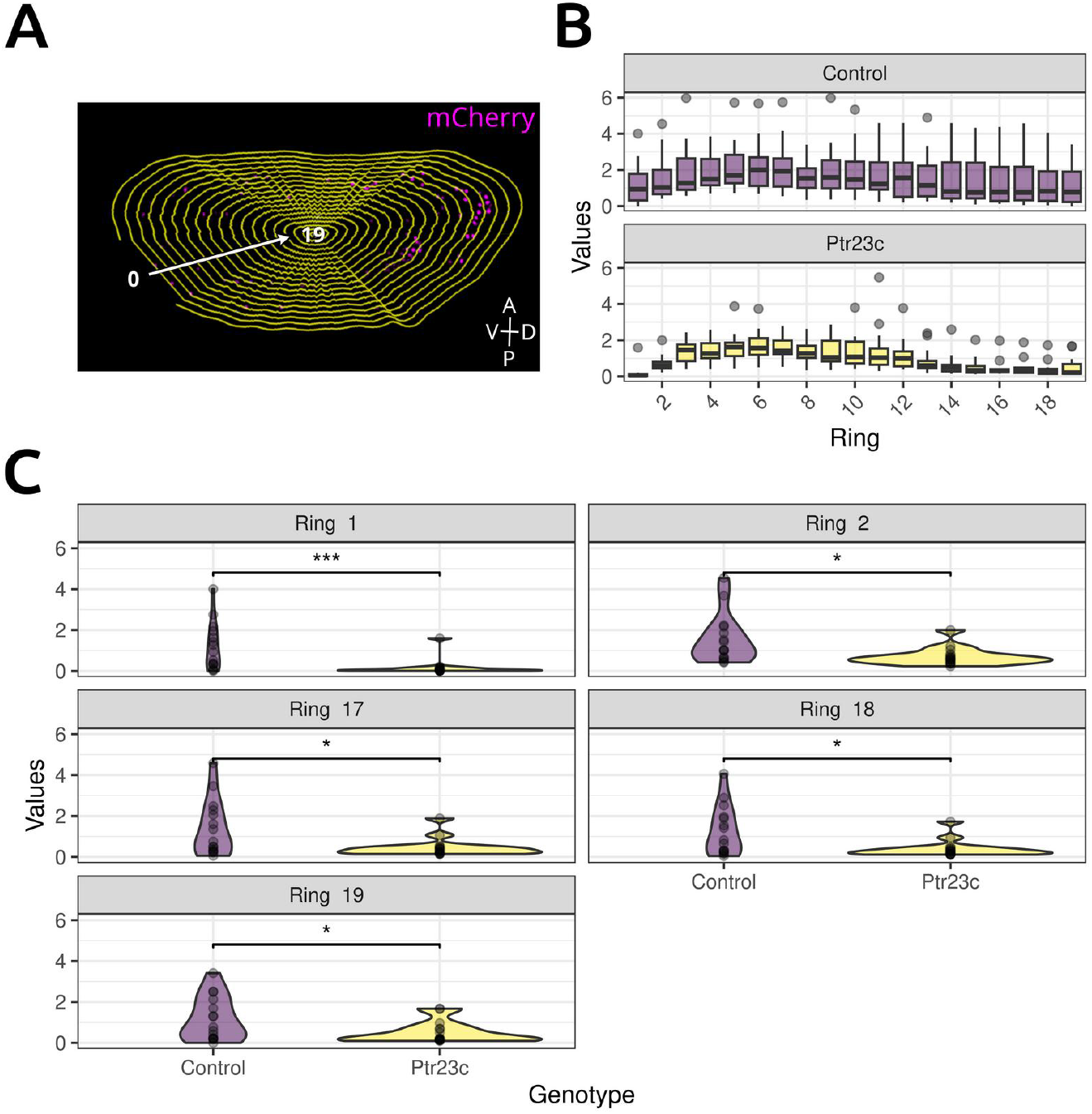
Hemocyte distribution at stage 16 in *Ptr*^*23c*^ embryos. (A) Integrated density of mCherry fluorescence detected in 19 concentric rings, defined as regions of interest (ROIs) numbered from the periphery (ring1) towards the center (ring 19). The two genotypes exhibited statistically different distributions (W = 53371, p = 8.613e-11, Wilcoxon rank sum test with continuity correction). (B) Rings with statistically significant differences are shown (rings 1, 2, 17, 18 and 19, Wilcoxon U-test, p<0.05). N = 15.

### Hemocyte proliferation and differentiation

Mitosis among embryonic hemocytes typically stops at stage 11 (Tepass et al., 1994). To investigate whether the observed hemocytes distribution at stage 16 in *Ptr*^*23c*^ mutants was the result of delayed early proliferation, we counted mitotic hemocytes at stage 10 using immunolabeling of mitotic mCherry-positive cells with antibodies against phosphorylated histone H3 (anti-PH3) (see Figure 3A). No statistical difference in the mitotic index was detected between *Ptr*^*23c*^ (median = 0.039, n = 8) and control (median = 0.011, n = 7) embryos at this stage (Figure 3B).

**Figure 3.**
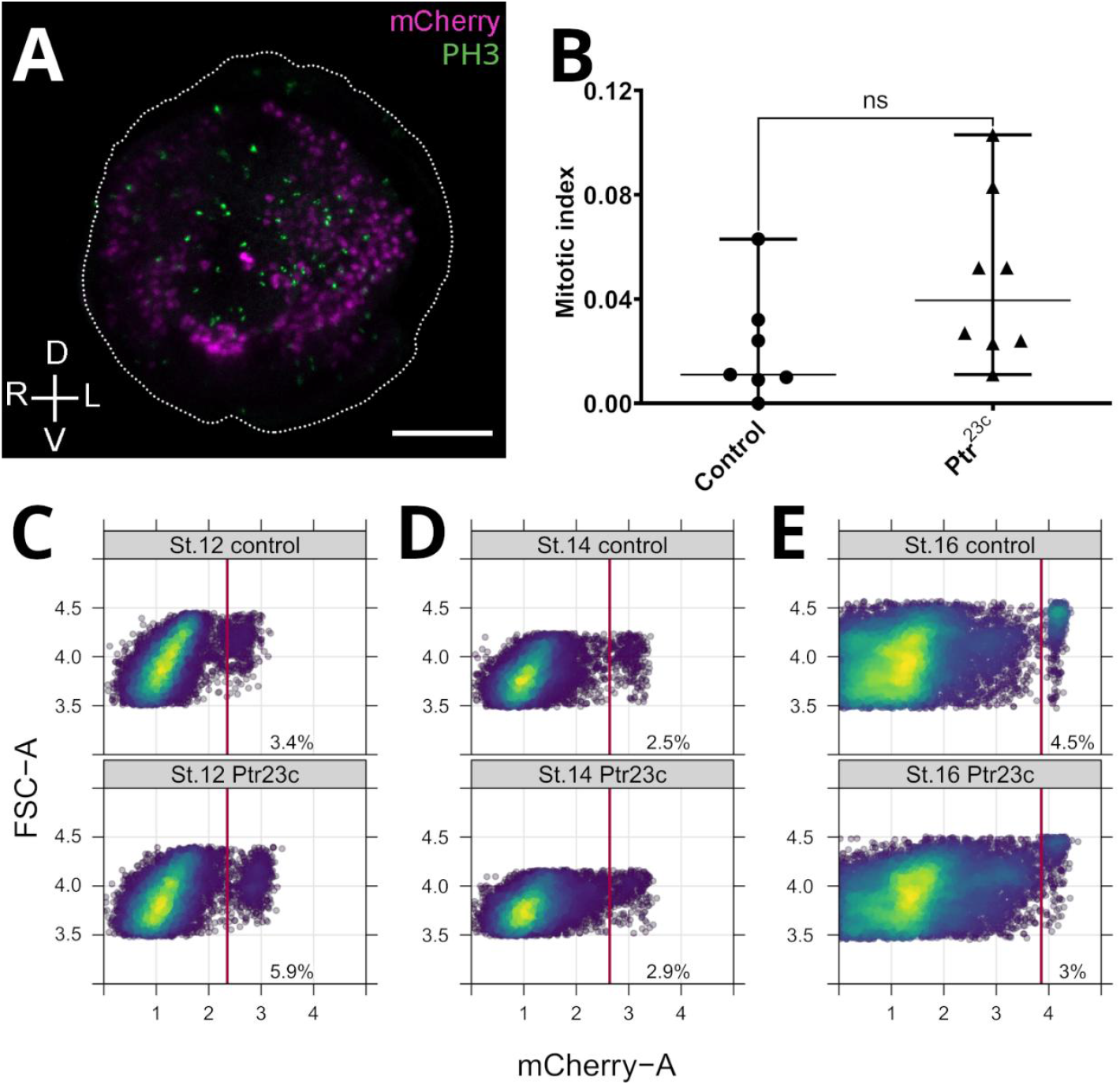
*Ptr*^*23c*^ mutants show a normal mitotic index and altered differentiation pattern in embryonic hemocytes. (A) Stage 10 embryos expressing nuclear mCherry were immunostained with an anti-PH3 antibody. The cephalic region of the embryo was imaged using confocal laser microscopy. (B) The number of double-positive nuclei was determined and the mitotic index was calculated. The medians were 0.0110 (control) and 0.0395 (*Ptr*^*23c*^) (Mann-Whitney U-test). (C-E) Time course of hemocyte differentiation in *Ptr*^*23c*^ mutants. Flow cytometry analysis of hemocyte population (*srp*::mCherry+ cells). (C) Based on cytometric evaluation of mCherry reporter fluorescence, a subset of hemocytes with higher fluorescence intensity was quantified. At stage 12 a greater number of hemocytes with high reporter expression were observed in *Ptr*^*23c*^ embryos than in control embryos. This represented a higher percentage (5.9%) of the total number of cells within the Non Debris/Singlets gate than in the control group (3.4%). (D) At stage 14, mCherry fluorescence increased in embryos, and the percentage difference between *Ptr*^*23c*^ hemocytes (2.9%) and control hemocytes (2.5%) decreased. (E) At stage 16, shortly after dispersal, hemocytes exhibited a peak in mCherry fluorescence and the percentage difference had reverted. Putative mature hemocytes from *Ptr*^*23c*^ embryos were 50% (3%) lower than in the control group (4.5%). N = 150. Scale bar: 50 μm. L: left, R: right, D: dorsal, V: ventral.

Flow cytometry analysis at stage 12, using *srp*Hemo-H2A::mCherry fluorescence as an indicator of differentiation, revealed that the fluorescence intensity of *srpHemo-H2A::mCherry*-positive hemocytes was 73.5% higher in *Ptr*^*23c*^ embryos (5.9%) than in controls (3.4%) (Figure 3C). This suggests that hemocytes in *Ptr*^*23c*^ embryos exhibit elevated *serpent* (*srp*) expression at stage 12.

By stage 14, both *Ptr*^*23c*^ and control embryos showed similar percentages of hemocytes with higher intensity *srpHemo-*H2A::mCherry fluorescence. This convergence suggests that hemocyte *srp* expression had stabilized by this stage (Figure 3D).

At stage 16, hemocytes reached their highest mCherry signal. However, *Ptr*^*23c*^ embryos exhibited 50% fewer hemocytes with high-intensity-fluorescence (3%) than controls (4.5%) (Figure 3E). Altered expression of the hemocyte differentiation marker *srp* could have significant implications for immune function and developmental viability (Shlyakhover et al., 2018; Bazzi et al., 2018; Monticelli et al., 2024).

## Discussion

In this study, we explored the impact of a null mutation (*Ptr*^*23c*^) of the gene *Ptr* in the regulation of the embryonic *Drosophila* hemocytes development. Our results suggest that *Ptr*^*23c*^ leads to significant disruptions in hemocyte distribution and the timing of differentiation, which could potentially affect the development of other tissues and immune processes (Bazzi et al., 2018; Monticelli et al., 2024).

Initially we hypothesized that *Ptr*^*23c*^ reduced the total number of hemocytes, because we had observed a decrease in hemocyte numbers in the ventral region of *Ptr*^*23c*^ mutant embryos at stage 16 (Bolatto et al., 2015). However, the present analysis, which is far more systematic and quantitative, revealed no significant differences in hemocyte counts. Similarly, the reduction in hemocytes could also not be explained by a shift in subpopulations towards CC differentiation. Interestingly, we also observed changes in the spatial distribution of hemocytes, specifically a reduction in their abundance in the peripheral and central regions of the *Ptr*^*23c*^ mutant embryo. This may explain the lower number of hemocytes previously observed in the ventral region (Bolatto et al., 2015).

Defects in hemocyte migration, as observed in *pvr* mutants, have been described to be associated with subsequent high hemocyte death by apoptosis (Brückner et al., 2004). Our quantification of the total number of apoptotic cells and of apoptotic hemocytes in *Ptr*^*23c*^ embryos revealed no increase relative to control embryos. This suggests the altered hemocyte distribution in the mutant was not due to cell loss or apoptosis-driven migration.

Interestingly, a reduction in the number of Repo-positive glia was observed in *Ptr*^*23c*^ embryos (Bolatto et al., 2022). This feature is also present in *repo* mutants, where reduced clearance of apoptotic bodies by phagocytic glia leads to secondary hemocyte recruitment. This represents a compensatory mechanism that counteracts glial loss by attracting hemocytes to the ventral nerve cord without altering the total hemocytes count (Armitage et al., 2020). Similarly, the changes in hemocyte distribution observed in *Ptr*^*23c*^ embryos may suggest a broader and homogeneous compensatory response to potential alterations in other developing tissues. This is consistent with our previous report of muscular defects in *Ptr*^*23c*^ larvae (Parada et al., 2025).

The hemocyte marker *srp*Hemo-H2A::3XmCherry not only allowed us to specifically detect hemocyte nuclei (Gyoergy et al., 2018) and the time at which the endogenous gene was switched on. Our flow cytometry analysis showed evident changes in the expression of the *srp*-directed fluorescent reporter in the *Ptr*^*23c*^ mutant, indicating an early onset or an accelerated differentiation of hemocytes. Srp is also required in the larval lymph gland during embryonic hematopoiesis, where it is necessary and sufficient to induce hemocyte development (Spahn et al., 2014). This suggests that the *Ptr*^*23c*^ mutation might accelerate the transition from progenitor cells to differentiated hemocytes.

Srp induces hemocyte differentiation by regulating the expression of phagocytic receptors (Shlyakhover et al., 2018), and the resulting plasmatocyte-mediated phagocytosis induces their own maturation (Weavers et al., 2016). Following their embryonic dispersal, hemocytes differentiate into mature cells while downregulating *srp* expression (Rehorn et al., 1996). The mechanisms governing this downregulation are not yet fully understood, but our observations suggest that the early differentiation of mCherry+ hemocytes in *Ptr*^*23c*^ mutants may cause premature *srp* downregulation. This could explain the reduced number of mCherry+ (and therefore probably mature) hemocytes observed in *Ptr*^*23c*^ mutants by stage 16. While a clear departure from the normal *srp* expression pattern is evident in the absence of *Ptr*, it would be desirable to further investigate this issue by using additional differentiation markers. Although early differentiation could affect hemocyte maturation, the observed changes in the distribution could be explained by compensatory mechanisms due to a reduced number of glial cells.

The molecular mechanisms underlying the premature differentiation of hemocytes in *Ptr*^*23c*^ mutants remain unclear. In the larva, hemocyte progenitors are kept quiescent or undifferentiated through via Hedgehog-Patched signaling (Csordás et al., 2021; Hultmark and Andó, 2022; Mandal et al., 2007; Tokusumi et al., 2018), which is activated by Srp (Tokusumi et al., 2010). Previous evidence suggested that Ptr acts as a negative regulator of the Hh pathway (Bolatto et al., 2022). Therefore, it is possible that Ptr affects other relevant signaling pathways through Hh signalling, such as those involving Notch or autophagy, which are known to regulate hemocyte differentiation and migration (Katz et al., 2024; Leitão and Sucena, 2015). Further investigation of these pathways is necessary to better understand how *Ptr* modulates hemocyte dynamics.

Our findings integrate Ptr into the complex signaling network governing hemocyte development in *Drosophila*. Analysis of the *Ptr*^*23c*^ mutant phenotype demonstrates that Ptr plays a key role in regulating hemocyte differentiation and spatial distribution.

In summary, our study identifies Ptr as a relevant factor in embryonic hemocyte development. Future studies should explore the precise molecular mechanisms by which Ptr influences hemocyte dynamics, as well as the broader consequences of the lack of Ptr deficiency on immune function and organismal development.

## Acknowledgements

The authors would like to thank Drs. Inés Carrera, Hugo Peluffo and Flavio Zolessi for their valuable advice and Daria Siekhaus for the fly stocks. They also gratefully acknowledge the Cell Biology Unit at the Institut Pasteur de Montevideo and Dr. María José Ferreiro from the Department of Experimental Neuropharmacology at the IIBCE for their support and assistance. Stock with hemocyte marker was kindly gifted by Dr. Daria Siekhaus was used in this study.

